# Abundant and rare bacterial taxa structuring differently in sediment and water in thermokarst lakes in the Yellow River Source area, Qinghai-Tibet Plateau

**DOI:** 10.1101/2021.05.26.445882

**Authors:** Ze Ren, Cheng Zhang, Xia Li, Kang Ma, Baoshan Cui

**Author notes:** **Authors contributed equally** (Ze Ren, Cheng Zhang). **Corresponding Author:** Xia Li, Advanced Institute of Natural Sciences, Beijing Normal University, Zhuhai 519087, China, Baoshan Cui, School of Environment, Beijing Normal University, Beijing 100875, China.

## Abstract

Thermokarst lakes are forming from permafrost thaw and severely affected by accelerating climate change. Sediment and water in these lakes are distinct habitats but closely connected. However, our understanding of the differences and linkages between sediment and water in thermokarst lakes remain largely unknow, especially from the perspective of community assembly mechanisms. Here, we examined bacterial communities in sediment and water in thermokarst lakes in the Yellow River Source area, Qinghai-Tibet Plateau. Bacterial taxa were divided to abundant and rare according to their relative abundance, and the Sorensen dissimilarity (β_sor_) was partitioned into turnover (β_turn_) and nestedness (β_nest_). The whole bacterial communities as well as the abundant and rare subcommunities differed substantially between sediment and water, in terms of taxonomical composition, α-diversity, and β-diversity. Sediment had significantly lower α-diversity indexes but higher β-diversity than water. Abundant taxa had significantly higher relative abundances but lower α-diversity and β-diversity than rare taxa. Moreover, bacterial communities are predominantly governed by strong turnover processes (β_turn_/β_sor_ ratio of 0.925). Abundant subcommunities were significantly lower in β_turn_/β_sor_ ratio compared to rare subcommunities. Bacterial communities in sediment had a significantly higher β_turn_/β_sor_ ratio than in water. The results suggest that the bacterial communities of thermokarst lakes, especially rare subcommunities or particularly in sediment, might be strongly structured by environmental filtering and geographical isolation, leading to compositional distinct. This integral study increased our current knowledge of thermokarst lakes, enhancing our understanding of the community assembly rules and ecosystem structures and processes of these rapid changing and vulnerable ecosystems.

## Introduction

Permafrost is widespread in high latitude and high elevation regions, covering about one quarter of land surface in the Northern Hemisphere and experiencing severe warming than the remainder of the globe (Qiu and Cheng, 1995; Zhang et al., 1999). Thermokarst lakes are forming as a result of permafrost thaw, acting as an important and widespread aquatic ecosystems in cold regions (Kokelj and Jorgenson, 2013; Farquharson et al., 2016) with significant roles in hydrological, ecological, and biogeochemical processes (Chin et al., 2016; In’T Zandt et al., 2020; Manasypov et al., 2021). In Arctic and sub-Arctic area, thermokarst lakes cover up to 40% of the permafrost area (de Jong et al., 2018). As an indicator of permafrost degradation, thermokarst lakes are suffering substantial changes in size and abundance owing to accelerating permafrost thaw (Karlsson et al., 2012; Pastick et al., 2019). Under accelerating climate change, the evolution process of thermokarst lakes including expansion, erosion, shrinkage, and disappearance, will be accelerated (Vincent et al., 2013; Luo et al., 2015; Biskaborn et al., 2019), resulting in significant impacts on regional environmental security and global biogeochemical processes (Luo et al., 2015). However, we are lacking of knowledge in ecosystem structure, function, and processes of thermokarst lakes compared to temperate lakes.

In lake ecosystems, bacteria play pivotal roles in ecosystem structuring and functioning. Bacterial communities exhibit high compositional and functional diversities and variabilities, with a relatively few abundant taxa coexist with considerable proportion of rare taxa (Lynch and Neufeld, 2015). In these enormously complex bacterial communities, abundant and rare taxa have fundamentally different characteristics and ecological roles (Logares et al., 2014). For example, abundant taxa contribute predominantly to biomass production and energy flow, whereas rara taxa contribute mostly to species richness and redundant functions (Pedros-Alio, 2012; Debroas et al., 2015). More and more studies have been conducted to unravel the differences between abundant and rare subcommunities in various environments, such as coastal and marine environments (Campbell et al., 2011; Logares et al., 2014), inland waters (Liu et al., 2015; Xue et al., 2018; Ren et al., 2020), and soil (Jiao and Lu, 2019; Xue et al., 2020). These studies indicated that abundant and rare subcommunities present contrasting community patterns and processes and are subject to distinct environmental factors. Assessment bacterial communities by considering abundant and rare subcommunities is also crucial for understanding the heterogeneous and fast changing thermokarst lake ecosystems.

In lake ecosystems, sediment and water are two distinct but closely interconnected environments (Carter et al., 2003; Parker et al., 2016). For example, these two environments have different chemical and physical properties but interact intimately through materials deposition from water to sediment and releasing from sediment to water (Roeske et al., 2012). These two habitats host different assemblages of microorganisms with tremendous diversity which play vital roles in maintaining and driving ecosystem structure and processes (Lozupone and Knight, 2007; Roeske et al., 2012; Ren et al., 2019). Moreover, the variation of bacterial communities in sediment and water are driven differently by a variety of factors (Gasol et al., 2002; Simek et al., 2008; Ren et al., 2019). For thermokarst lakes, their formation (permafrost thaw) and evolution (horizontal and vertical permafrost degradation) mechanisms suggest that sediment and water have very close relationships in thermokarst lakes. Thermokarst processes stimulate the release of carbon, nutrients, and even heavy metals from deeper permafrost (sediment) to water and this releasing processes can be further accelerated by microbial activities and climate change (Bowden, 2010; In’T Zandt et al., 2020; Manasypov et al., 2021). Intensifying climate change is beginning to unlock more materials and microorganisms from sediment to water (Graham et al., 2012; Mackelprang et al., 2017). In cold region, thermokarst lakes are especially important due to their huge abundance (Polishchuk et al., 2017), massive storage of water, carbon, and nutrients (Reyes and Lougheed, 2015), as well as their enormous contributions of greenhouse gases (Walter et al., 2006; Serikova et al., 2019; In’T Zandt et al., 2020). Although immense amounts of microbial research have been conducted in thermokarst lakes, overwhelming majority of which focused on surface water (TRANVIK, 1989; Shirokova et al., 2013; Vucic et al., 2020) rather than sediment or both. However, the differences and linkages between sediment and water in thermokarst lakes are remain largely unknow.

As the third pole of the world, Qinghai-Tibet Plateau is covered by permafrost up to 40% of the land area and is highly sensitive to climate change (Zou et al., 2017). The ongoing global warming has accelerated permafrost degradation, resulting in extensive changes of thermokarst lakes with lake number increase and lake area expansion (Luo et al., 2015; Zhang et al., 2017). In this study, we investigated the bacterial communities in sediment and water in thermokarst lakes of the Yellow River Source area. Our object was to reveal

1. the assemblage structure processes of abundant and rare taxa in sediment and water,
2. the responses of abundant and rare subcommunities to environmental variables, and
3. the differences and linkages between sediment and water. The integral understanding of bacterial community in both sediment and water could provide insights into community assembly rules and ecosystem structures and processes of the thermokarst lakes.

## Methods

### Study area, field sampling, and chemical analysis

This study was conducted in the Yellow River Source area on the Qinghai-Tibet Plateau (Figure 1). In early July 2020, we sampled 23 thermokarst lakes in the study area. The elevation of the studied lake surface ranged from 4200 to 4350 m. In each lake, both water samples and sediment samples were collected. Conductivity and pH of the lake water were measured *in situ* using a multiparameter instrument (YSI ProPlus, Yellow Springs, Ohio). Because the thermokarst lakes are very shallow, only surface water samples were collected at the depth of 0.3 to 0.5m. Three 1L water samples were filled in acid clean bottles and transported to the laboratory for chemical analyses, including dissolved organic carbon (DOC), total nitrogen (TN), and total phosphorus (TP). Microbial samples were collected by filtering 200 mL water onto a 0.2 μm polycarbonate membrane filter (Whatman, UK) and frozen in liquid nitrogen immediately in the field and stored at −80 □ in the lab until DNA extraction. Sediment samples were collected using a Ponar Grab sampler. The top 5 cm of the sediment was collected and homogenized. Sediment microbial samples were connected in a 45 mL sterile centrifuge tube and frozen in liquid nitrogen in the field and stored at −80 □ in the lab for DNA extraction. The remaining sediments were air-dried for the determination of chemical properties, including pH, conductivity, sediment organic carbon (SOC), TN, and TP. The basic chemical properties of sediment and water samples were summarized in Table S1.

**Figure 1.**
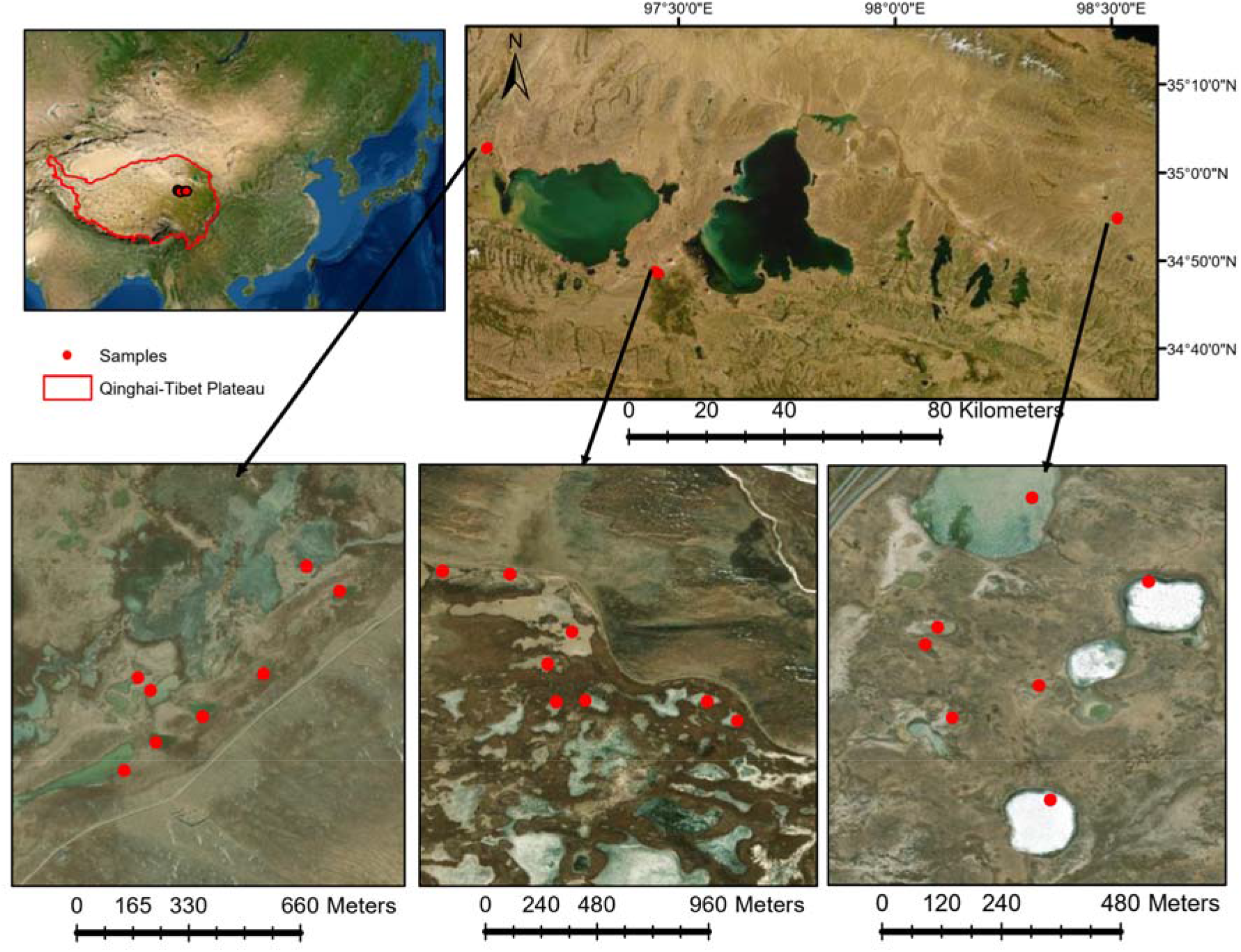
Water and sediment samples were collected from 23 lakes in early July 2020 in the Yellow River Source area on the Qinghai-Tibet Plateau.

### DNA extraction, PCR, and sequencing

The water (filter) and sediment (0.5g) microbial samples were used to extract genomic DNA using the DNeasy PowerSoil Kit (QIAGEN, Germany) following the manufacturer’s protocols. The genome DNA was used as template for PCR amplification with the barcoded primers and Tks Gflex DNA Polymerase (Takara, USA). The hypervariable V3-V4 regions of bacterial 16S rRNA were amplified using universal primers 343F 5’-TACGGRAGGCAGCAG-3’ and 798R 5’-AGGGTATCTAATCCT-3’ (Nossa et al., 2010). In order to minimize amplification bias, three individual PCR amplifications were performed using the following procedure: initial denaturation at 94 °C for 5 min, 24 cycles of denaturation at 94 °C for 30 s followed by annealing at 56 °C for 30 s and extension at 72 °C for 20 s, and final extension step at 72 °C for 5 min. Amplified DNA was verified by agarose gel electrophoresis, purified using the AMPure XP beads (Beckman, USA), and quantified using Qubit dsDNA assay kit (Thermo Fisher Scientific, USA). Sequencing of the amplicon libraries was conducted on an Illumina MiSeq platform (Illumina, San Diego, CA, USA) according to manufacturer’s instructions. Raw sequence data can be accessed at the China National Center for Bioinformation (CRA004269 under the project PRJCA005279).

### Data analyses

Raw sequence data were preprocessed using Trimmomatic software (version 0.35) (Bolger et al., 2014) to detect and cut off ambiguous bases and low-quality sequences with average quality score below 20. After trimming, paired-end reads were assembled using FLASH software (Reyon et al., 2012). Parameters of assembly were: 10bp of minimal overlapping, 200bp of maximum overlapping and 20% of maximum mismatch rate. Sequences were performed further denoising using QIIME 1.9.1 (Caporaso et al., 2010) as follows: reads with ambiguous, homologous sequences or below 200bp were abandoned; reads with 75% of bases above Q20 were retained; reads with chimera were detected and removed. Clean reads were subjected to primer sequences removal and clustering to generate operational taxonomic units (OTUs) against the SILVA 132 database (Quast et al., 2013) using QIIME. To avoid the bias of surveying efforts, the sequence data were normalized at the depth of 27,890 sequence per sample (Figure S1).

Bacterial taxa were defined as abundant and rare according to their relative abundance (Pedros-Alio, 2012; Logares et al., 2014). In this study, OTUs with a relative abundance ≥ 0.1% of the total sequences were defined as abundant and OTUs with a relative abundance <0.01% were defined as rare. The α-diversity indices, including Chao 1, observed OTUs, Shanon, and phylogenetic diversity (PD whole tree) were calculated using QIIME 1.9.1 (Caporaso et al., 2010). To further reveal the mechanisms underlying the discrepancies observed between the abundant and rare subcommunities, as well as between bacterial communities in sediment and water, the Sorensen dissimilarity (β_sor_, a β-diversity metric that measures compositional differences between sites) was partitioned to turnover component (β_turn_) and nestedness□resultant fraction (β_nest_) (Baselga, 2010) using the “betapart 1.5.4” package (Baselga and Orme, 2012). The differences of whole, abundant, and rare communities/sub-communities between sediment and water were revealed by principal coordinates analysis (PCoA) based on Bray-Curtis distance and tested by ADONIS (analysis of variance using distance matrices), ANOSIM (analysis of similarity), and MRPP (multi-response permutation procedure analysis) using “vegan 2.5-7” package (Oksanen et al., 2007). Structural equation model (SEM) was conducted using “piecewiseSEM 2.1.2” package (Lefcheck, 2016) to quantify the effects of environmental variables on β-diversity of bacterial communities in sediment and water, as well as the biological and physicochemical relationships between sediment and water. All the analyses were conducted in R 4.0.4 (R Core Team, 2017).

## Results

### Bacterial communities in sediment and water

After quality filtering and the removal of chimeric sequences, 1,282,940 high-quality sequences were acquired and clustered into 27,889 OTUs at 97% nucleotide similarity level in sediment and water of the studied thermokarst lakes. In total, 17178 OTUs were detected in sediment and 17455 OTUs were detected in water. For the assigned OTUs in sediment, Proteobacteria (30%), Bacteroidetes (29%), Firmicutes (28%), and Actinobacteria (6%) were the dominant phyla (with an average relative abundance >5%, Figure 2a). In water bacterial communities, Proteobacteria (40%), Bacteroidetes (30%), Actinobacteria (11%), and Acidobacteria (5%) were the dominant phyla (Figure 2a). In terms of the top 10 most abundant phyla in these thermokarst lakes, sediment had significantly higher Firmicutes and Spirochaetes, while lower Acidobacteria, Actinobacteria, Fusobacteria, Nitrospirae, Patescibacteria, and Proteobacteria than water (Figure 2a). Moreover, bacterial α-diversity was significantly lower in sediment than in water in terms of Chao 1, observed OTUs, and phylogenetic diversity (Figure 2b).

**Figure 2.**
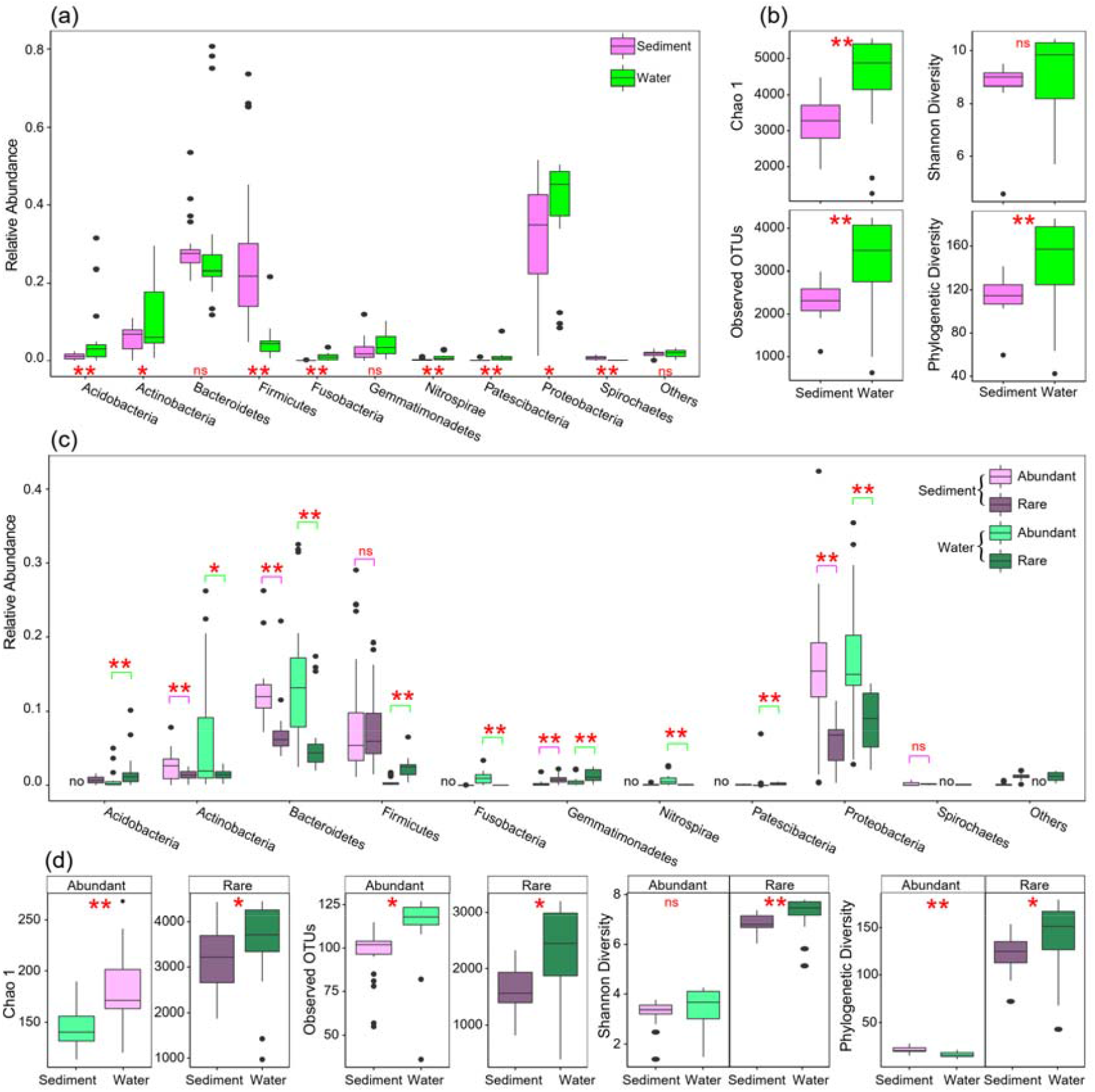
Bacterial community composition and alpha diver in sediment and water of the studied thermokarst lakes. (a) Differences in taxonomic composition of overall communities between sediment and water. (b) Differences in α-diversity of overall communities between sediment and water. (c) Differences in taxonomic composition between abundant and rare subcommunities. (d) Differences in α-diversity between abundant and rare subcommunities. The differences were assessed using Wilcoxon rank-sum test with “ns”, “*”, and “**” represent non-significance, P<0.05, and P<0.01, respectively.

Venn diagram shown that sediment and water harbored a large amount of unique OTUs which were only presented in sediment (n=10434) or water (n=10711) while only 6744 OTUs were detected in both (Figure 3a). Substantial differences were observed between sediment and water bacterial communities based on the PCoA results (Figure 3b). These results were further confirmed by the results of adonis, ANOSIM, and MRPP analyses (Table S2). The sediment bacterial communities had a significantly higher β-diversity (Figure 3c) but lower niche width (Figure 3d) than water ones. The results indicate that the sediment and water environments harbored distinct bacterial communities in terms of taxonomic composition as well as α- and β-diversity.

**Figure 3.**
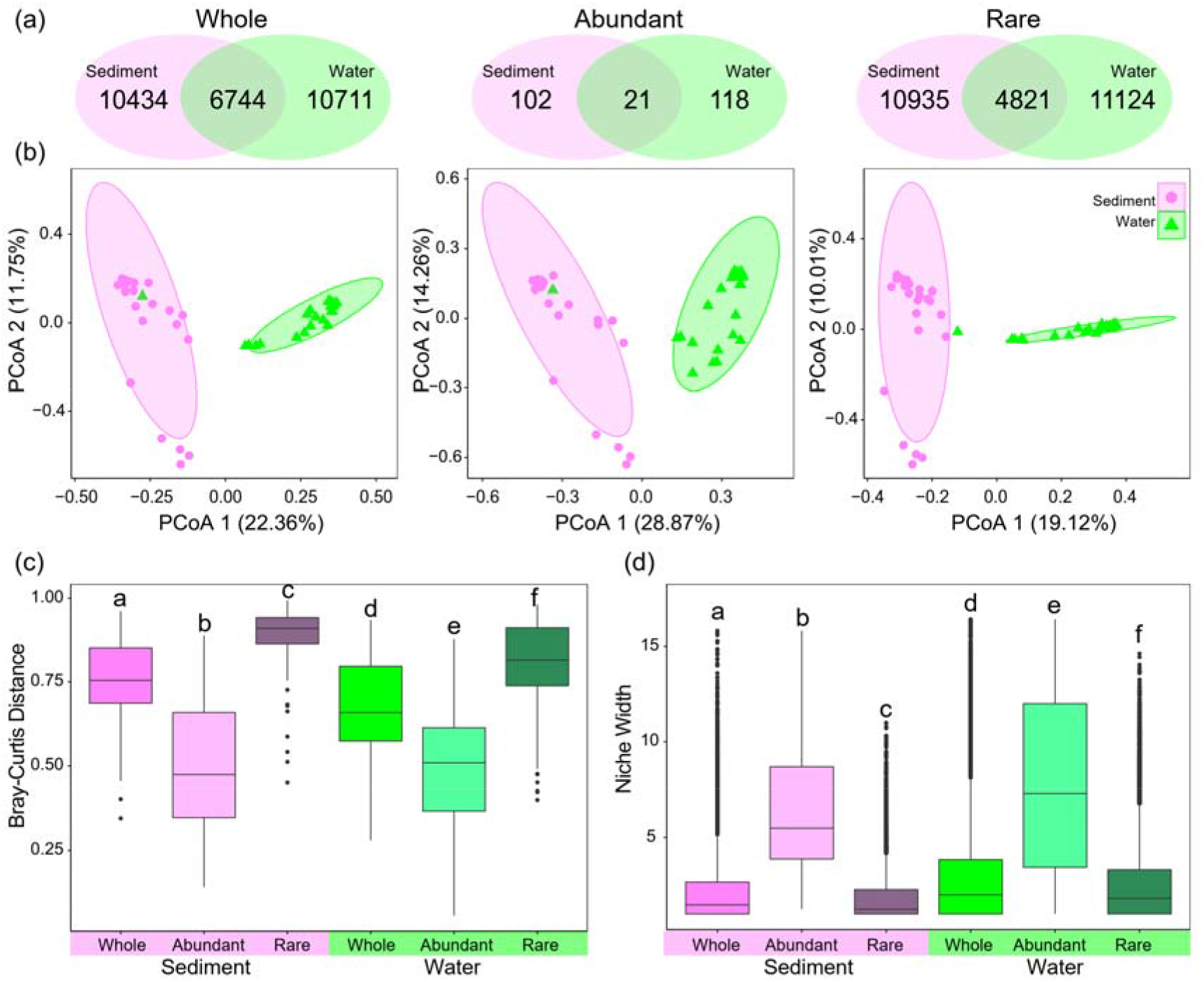
Structural differences of bacterial communities between sediment and water in terms of whole, abundant, and rare subcommunities. (a) Venn diagram showing the unique and shared OTUs in sediment and water. (b) Principal coordinates analysis (PCoA) based on Bray-Curtis distance using the relative abundance of OTUs. (c) β-diversity measured using Bray-Curtis distance. (d) Niche width of the OTUs. The different low-case letters in panels (c) and (d) represent significant difference assessed using ANOVA.

### Abundant and rare subcommunities in sediment and water

In the whole bacterial communities, a large proportion of the OTUs were identified as rare taxa (69.6% for sediments and 69.8% for water in terms of the number of OTUs), but they only accounted for 25.1% and 22.3% of relative abundance in sediments and water (Figure S2), respectively. Conversely, a very small proportion of the OTUs were identified as abundant taxa (4.2% and 3.9% in both sediments and water, respectively), which account for 39.8% and 40.7% of the average relative abundance in each sample (Figure S2). Venn diagram shown that only 21 abundant OTUs and 4821 rare OTUs were abundant in both sediment and water environments (Figure 3a).

Abundant subcommunities were substantially different in taxonomic composition comparing with rare subcommunities in both sediment and water (Figure 2c). In sediment, abundant subcommunities had significantly higher Actinobacteria, Bacteroidetes, Firmicutes, and Proteobacteria but lower Gemmatimonadetes than rare subcommunities (Figure 2c). In water, abundant subcommunities had higher Actinobacteria, Bacteroidetes, Fusobacteria, Nitrospirae, Patescibacteria, and Proteobacteria but lower Acidobacteria, Firmicutes, and Gemmatimonadetes than rare subcommunities (Figure 2c).

Moreover, abundant and rare subcommunities also had distinct structure between sediment and water (Figure 3b). When comparing abundant and rare subcommunities, abundant subcommunities had a significantly lower β-diversity (Bray-Curtis distance) but higher niche width than rare subcommunities in both sediment and water (Figure 3c, d). When comparing sediment and water, abundant subcommunities had lower β-diversity and niche width in sediment than in water, while rare subcommunities had higher β-diversity but lower niche width in sediment than in water (Figure 3c, d). The results indicate that, there were distinct distribution patterns and taxonomic composition between abundant and rare subcommunities in sediment and water.

### Turnover and nestedness

Based on beta-partitioning, the Sorensen dissimilarity (β_sor_, a β-diversity metric that measures compositional differences between sites) was partitioned to turnover component (β_turn_) and nestedness□resultant fraction (β_nest_) (Figure 4a). The estimated β_sor_ of the whole bacterial communities was 0.733±0.1, 0.827±0.04, and 0.638±0.14 for paired sites in terms of only sediment samples, between sediment and water samples, and only water samples, respectively (Figure 4a). The estimated β_sor_ of abundant subcommunities was 0.273±0.1, 0.365±0.08, and 0.198±0.11 for paired sites in terms of only sediment samples, between sediment and water samples, and only water samples, respectively (Figure 4a). The estimated β_sor_ of rare subcommunities was 0.786±0.09, 0.871±0.04, and 0.691±0.14 for paired sites in terms of only sediment samples, between sediment and water samples, and only water samples, respectively (Figure 4a). Abundant subcommunities had a significantly lower β_sor_ than rare subcommunities (Figure 4a). Sediment communities had a significantly higher β_sor_ than water communities (Figure 4a).

**Figure 4.**
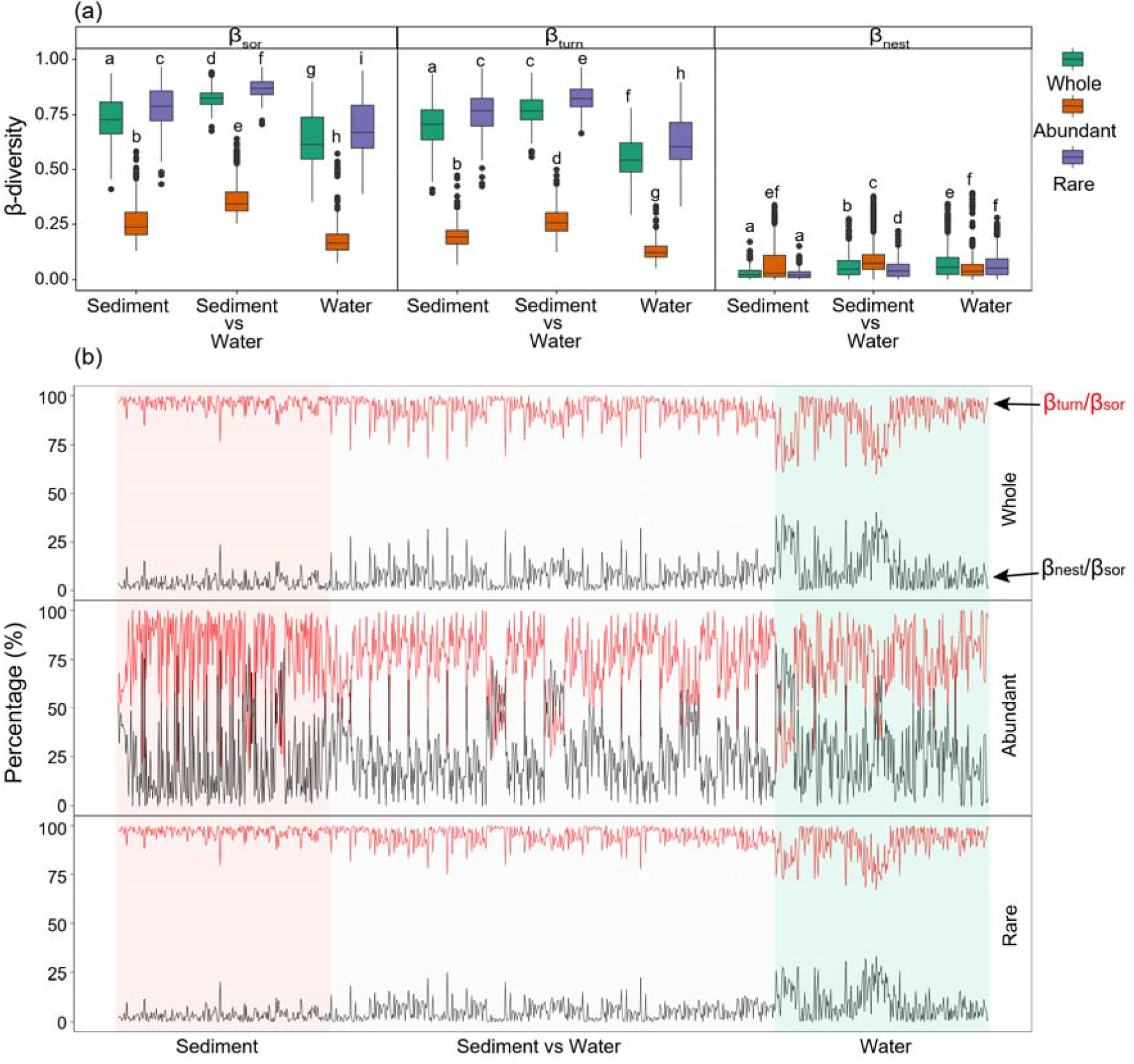
Beta-diversity partitioning results between pairs of sites in terms of only sediment samples, between sediment and water samples (sediment vs water), and only water samples. (a) Differences in total β-diversity calculated as the Sorensen dissimilarity (β_sor_), turnover component (β_turn_), and nestedness□resultant fraction (β_nest_) between the whole, abundant, and rare assemblages. The different low-case letters represent significant difference assessed using ANOVA. (b) The contributions of turnover component to total Sorensen dissimilarity (β_turn_/β_sor_ ratio) and of nestedness□resultant fraction to total Sorensen dissimilarity (β_nest_/β_sor_ ratio) for paired sites.

Moreover, the estimated β_sor_ of bacterial communities was mainly contributed by turnover component with an average β_turn_/β_sor_ ratio of 0.925 for the whole communities (Figure 4b). Abundant subcommunities had a significantly lower contribution of β_turn_ to β_sor_ (β_turn_/β_sor_ ratio of 0.746) than that of rare subcommunities (β_turn_/β_sor_ ratio of 0.942) (Figure 4b and Figure S3). Comparing different habitats, sediment had a significantly higher β_turn_/β_sor_ ratio than water (Figure 4b and Figure S3).

### Environmental responses of abundant and rare subcommunities in sediment and water

The results of SEM revealed the relationships between the variations of environmental variables and the variations of bacterial communities, as well as the relationships between sediment and water (Figure 5). Based on the SEM results, sediment and water had close associations in conductivity and total phosphorus (Figure 5). In addition, sediment and water had close associations in terms of the β-diversity of the whole bacterial communities as well as the abundant subcommunities (Figure 5). In sediment, pH and TP had positive effects on the β-diversity of the whole communities, TP had positive effects on β-diversity of the abundant subcommunities, and pH had positive effects on β-diversity of the rare subcommunities (Figure 5). In water, however, conductivity had negative effects, and TN and TP had positive effects on the β-diversity of the whole communities as well as the abundant and rare subcommunities (Figure 5).

**Figure 5.**
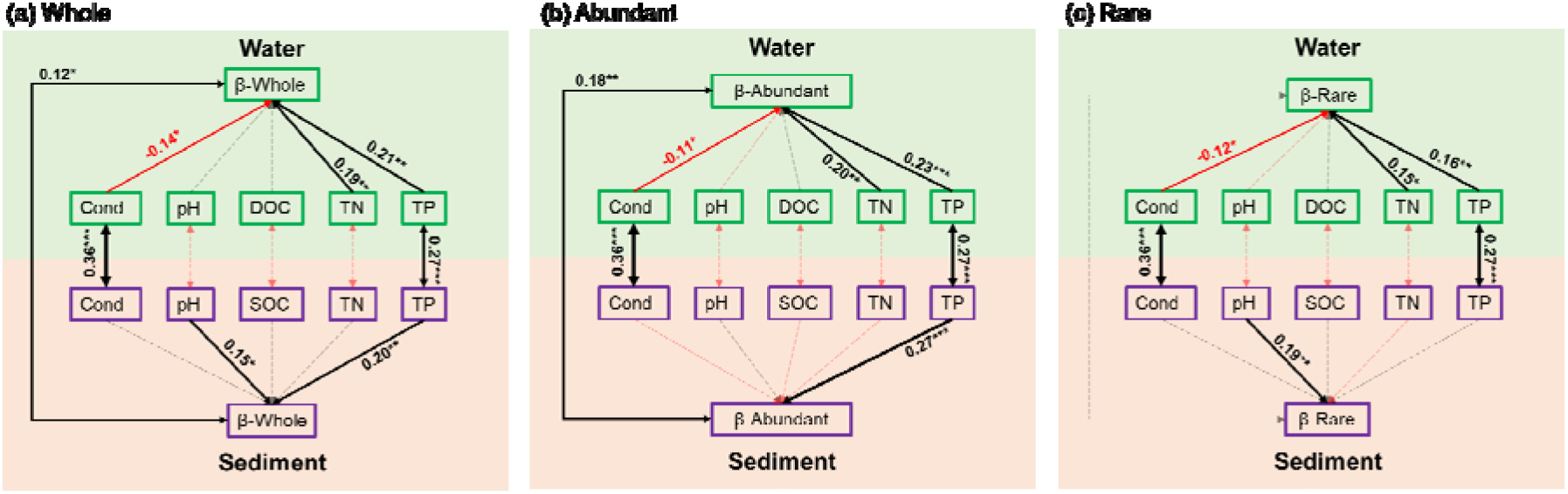
Structural equation model illustrating the relationships between the variation of environmental variables and the β-diversity of bacterial communities in sediment and water in terms of (a) whole community, (b) abundant subcommunities, and (c) rare subcommunities. Solid and dashed arrows represent the significant and nonsignificant relationships, respectively. Red and black arrows represent negative and positive relationships, respectively. The significant path coefficients were shown adjacent to the path with *, **, and *** denote the significant level of p<0.05, p<0.01, and p<0.001, respectively.

For different components of β-diversity, pH and TP had positive effects on β_turn_ of the whole communities and rare subcommunities in sediment (Figure 6). In water, however, environmental variables had more significant effects (Figure 6). For the whole water bacterial communities, TP had positive effects on β_turn_, conductivity had negative while TN and TP had positive effects on β_nest_ (Figure 6). For the abundant subcommunities in water, TN and TP had positive effects on β_turn_, conductivity and pH had negative while TN and TP had positive effects on β_nest_ (Figure 6). For the rare subcommunities in water, TN and TP had positive effects on β_turn_, conductivity had negative while TN had positive effects on β_nest_ (Figure 6).

**Figure 6.**
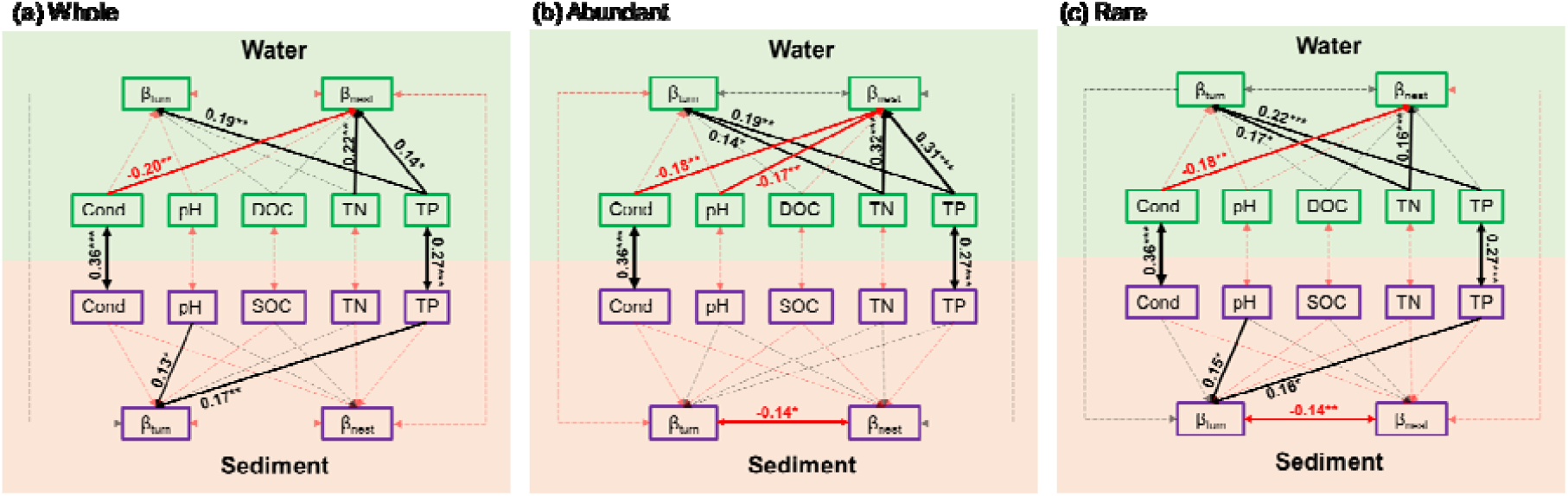
Structural equation model illustrating the relationships between the variation of environmental variables and different components of β-diversity (turnover component, βturn and nestedness□resultant fraction, βnest) in sediment and water in terms of (a) whole community, (b) abundant subcommunities, and (c) rare subcommunities. Solid and dashed arrows represent the significant and nonsignificant relationships, respectively. Red and black arrows represent negative and positive relationships, respectively. The significant path coefficients were shown adjacent to the path with *, **, and *** denote the significant level of p<0.05, p<0.01, and p<0.001, respectively.

## Discussion

### Differences and linkages between sediment and water

This study showed that the whole bacterial communities as well as the abundant and rare subcommunities differed substantially between sediment and water, in terms of taxonomical composition, α-diversity, and β-diversity. It is unequivocal that different habitats usually harbor distinct microbial assemblages (Fierer et al., 2012; Hugerth et al., 2015; Louca et al., 2016). Previous studies have demonstrated that, in lake ecosystems, sediment and water host different bacterial communities (Gough and Stahl, 2011; Yang et al., 2016; Ren et al., 2019). In our study, eight out of the ten most abundant phylum presented significantly different relative abundance between sediment and water (Figure 2a). Specifically, Firmicutes had a mean relative abundance of 28% in sediment but only 5% in water and overwhelming majority of the Firmicutes OTUs were affiliated to Clostridia class, which are anaerobic and always dominate the bacterial communities during decomposition process in lake sediments (Xing et al., 2011; Zhao et al., 2017). In general, sediments always have high species-level diversity than water, especially in temperature lakes (Lozupone and Knight, 2007; Ren et al., 2019). In our study, however, sediment had significantly lower α-diversity indexes than water (Figure 2b) suggesting that the water column provides more niches for bacterial taxa than sediment. This was further supported by that the bacterial taxa had higher niche widths in water than in sediment (Figure 3d).

In addition, taxonomic composition, α-diversity, and β-diversity also differed substantially between abundant and rare subcommunities. Abundant and rare taxa may have distinct intrinsic characteristics and life strategies to survive in challenging environments (Logares et al., 2014; Malik et al., 2020). In bacterial communities, abundant taxa play as the core species with high growth rates and enormous contributions to most ecosystem functions (Pedros-Alio, 2006). On the contrary, rare taxa encompass ecologically redundant taxa that have low growth rates and competition capability, maintain community stability because of high redundancy, and contribute to community dissimilarity because of broad diversity (Pedros-Alio, 2012). There is no doubt that the abundant taxa had significantly higher relative abundances but lower α-diversity than rare taxa (Figure 2). In our studied bacterial communities, only 4% OTUs were abundant taxa which accounted for 40% of sequences, while more than 70% OTUs were rare taxa which only account for 24% of sequences (Figure S2). Moreover, in line with previous studies in many other ecosystems (Wu et al., 2016; Ren et al., 2020; Shu et al., 2021), abundant taxa had significantly lower β-diversity and occupy higher niche width than rare taxa (Figure 3), suggesting that abundant taxa were more ubiquitous distributed and more competitive for limited resources than rare taxa during the adaptation process (Malik et al., 2020).

In aquatic ecosystems, bacterial communities are controlled differently by various factors and processes in different environments (Gasol et al., 2002; Simek et al., 2008; Ren et al., 2019). To understand the factors that control bacterial community variations is a central theme in ecology (Fierer and Jackson, 2006; Pla-Rabes et al., 2011). In our study, the variations of bacterial communities had close relationships with pH and TP in sediment while with conductivity, TN, and TP in water (Figure 5). Moreover, the abundant and rare subcommunities respond different to environmental variables in sediment that abundant subcommunities only had a close relationship with TP while rare subcommunities only had a close relationships with pH (Figure 5). The availabilities of key nutrients (N and P) have long been demonstrated to be essential in structuring bacterial communities (Torsvik et al., 2002; Lee et al., 2017). Nutrient dynamics usually have close interactions with bacterial communities in water and sediment in lake ecosystems (Lee et al., 2017; Ren et al., 2019). The different responses of microorganisms to nutrients root in their metabolic features and ecological strategies, as well as environmental properties (Carbonero et al., 2014). Salinity is always the major environmental determinant of microbial community in aquatic environments (Lozupone and Knight, 2007). In thermokarst lakes, sediment was formed from permafrost soil. Many studies have demonstrated that soil bacterial communities are strongly affected by pH (Fierer et al., 2012; Ren et al., 2021).

In addition to the differences, sediment and water also had close biological and physicochemical relationships (Roeske et al., 2012; Parker et al., 2016). In our study, sediment and water were closely related in conductivity and TP (Figure 5). Moreover, the β-diversity, especially the β-diversity of abundant subcommunities, also had a significant relationship between sediment and water (Figure 5). These connections between sediment and water might be the results of thermokarst lakes formation and evolution processes, during which, nutrients and other elements are continuously released to water from permafrost and sediment (Mackelprang et al., 2017; In’T Zandt et al., 2020; Manasypov et al., 2021).

### Beta diversity and its different components

Unravelling the underlying mechanisms driving species distribution patterns is a vital issue in ecology and biogeography (Podani and Schmera, 2016). As a key term for assessing spatial and temporal variations of microbial assembly, β-diversity can be decomposed into two distinct components: turnover and nestedness (Baselga, 2010; Legendre, 2014), which may reflect the relative importance of different underlying mechanisms in structuring communities varies with the spatiotemporal scales (Podani and Schmera, 2016; Viana et al., 2016). In our study, bacterial communities are predominantly governed by strong turnover processes (β_turn_/β_sor_ ratio of 0.925 for the whole communities). However, abundant subcommunities were significantly lower in β_turn_/β_sor_ ratio compared to rare subcommunities (Figure 4b and Figure S3). Moreover, bacterial communities in sediment had a significantly higher β_turn_/β_sor_ ratio than in water (Figure 4b and Figure S3). Species turnover refers to the fact that species experiencing replacement of each other (gains and losses) along ecological gradients as a consequence of spatiotemporal constraints and/or environmental sorting (Leprieur et al., 2011). Therefore, we could expect that regions have high species turnover would also possess great heterogeneities in contemporary environmental conditions and/or strong geographical isolation caused by dispersal barriers (Gaston et al., 2007a, 2007b). In our studied thermokarst lakes, the environmental factors had high variations in both sediment and water (Table S1). However, sediment is more isolated than water between lakes, even for the intimate neighbors in the space. Thus, it is expected that sediment had higher β_turn_/β_sor_ ratio than water. Moreover, the ecological tolerance and niche breadth of taxa are also determinative factors for turnover rate (Leprieur et al., 2011), which is the potential reason that abundant taxa had lower β_turn_/β_sor_ ratio than rare taxa because of higher niche width and ecological tolerance of abundant taxa. In contrast to turnover, nestedness implies another pattern of richness difference that the species at a depauperate site is a subset of assemblages of a species-rich site (Leprieur et al., 2011; Baselga, 2012). The nestedness richness difference (species loss or gain) may result from various ecological processes, such as nestedness of habitats, selective colonization and/or extinction, as well as interspecific variation of environmental tolerance (Whittaker and Fernandez-Palacios, 2007; Baselga, 2010). All in all, our results suggest that the driving mechanisms for the variation of bacterial assemblages are differed in term of different habitats or relative abundances of taxa. The bacterial communities of thermokarst lakes, especially rare subcommunities or particularly in sediment, might be strongly structured by environmental filtering and geographical isolation which potentially driving these assemblages tend to be more compositionally distinct.

Integrating the differences in taxonomic composition and β-diversity patterns of bacterial communities in sediment and water in terms of the whole, abundant, and rare communities/sub-communities, we can propose a hypothesis that the bacterial communities in water and sediment are initially originated from the same source (permafrost) and then diverged into two distinct communities due to the environmental divergence as well as the different assemblage rules in constructing communities. The further study of the bacterial community divergence during the thermokarst lakes formation and evolution processes would promote our understanding of the ecological consequences of future climate change.

## Conclusions

Thermokarst lakes are among the world’s most pristine aquatic ecosystems and extremely vulnerable to accelerating climate change. Assessing the community composition and spatial pattern, as well as their underlying driving mechanisms in sediment and water is pivotal in understanding the heterogeneous and fast changing thermokarst lake ecosystems. This study highlights the differences and linkages of sediment and water in terms of physicochemical properties, taxonomical composition, and diversity patterns. Moreover, the underlying mechanisms driving taxa distributions patterns were also be revealed. There were distinct β-diversity patterns existed between abundant and rare subcommunities as well as between sediment and water. This integral study of bacterial communities can enhance our understanding of the community assembly rules and ecosystem structures and processes of the thermokarst lakes.

## Acknowledgements

We are grateful to the anonymous reviewers for the comments, to the Administration Office of Sanjiangyuan National Park for the assistances in the field work.

## Supplementary Information

**Figure S1.**
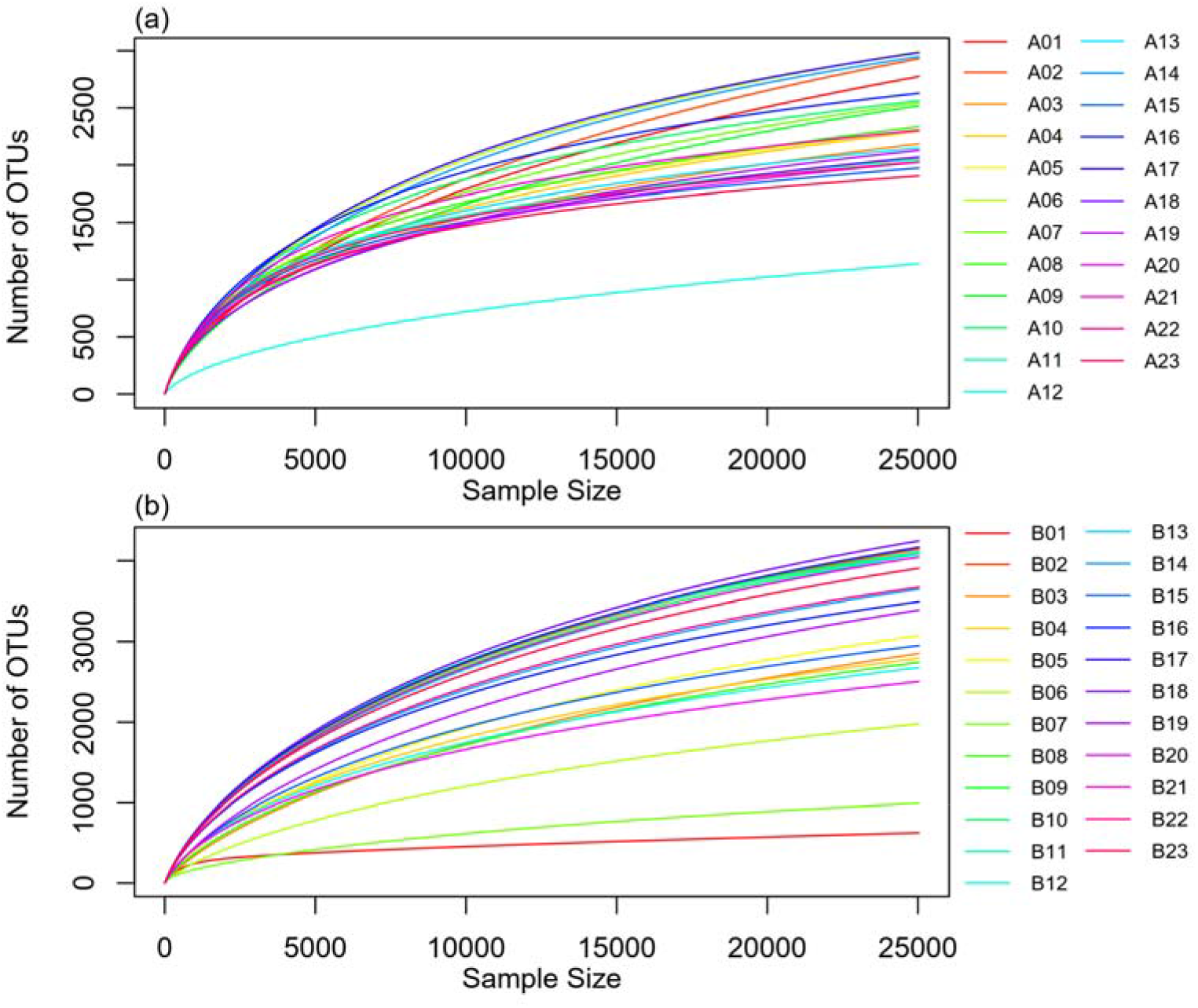
Rarefaction curve of operational taxonomic units (OTUs) in each sample site at 97% nucleotide sequence identity threshold. (a) sediment samples. (b) water samples.

**Figure S2.**
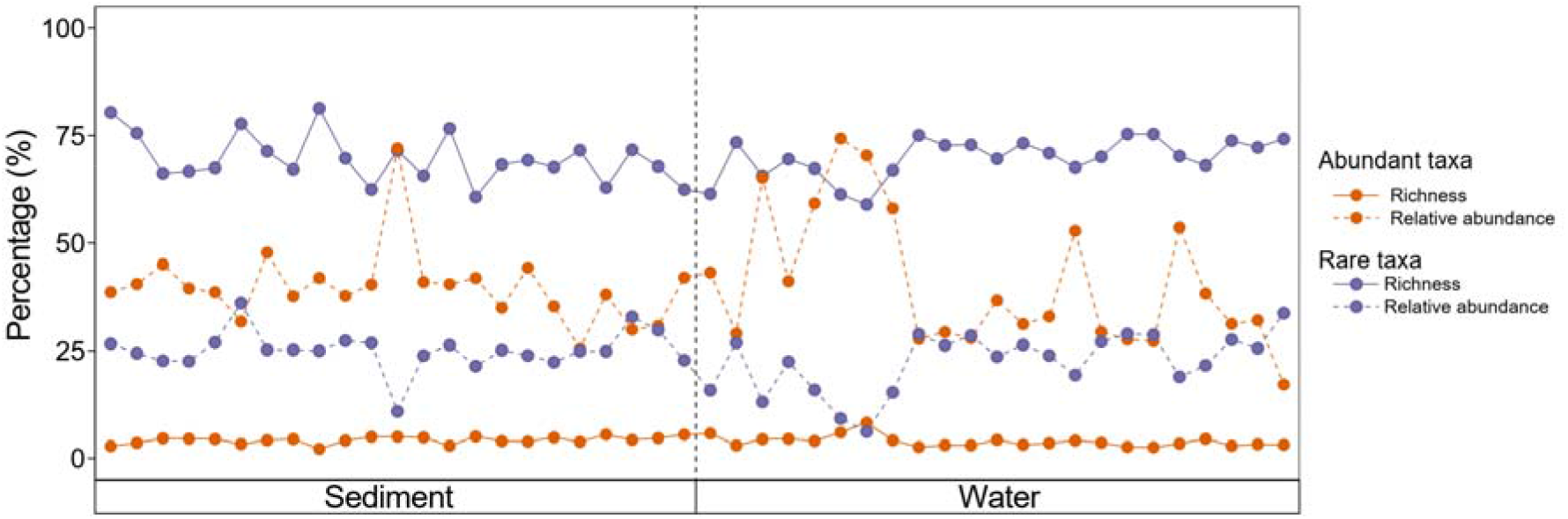
The proportion of the richness and relative abundance of OTUs in abundant and rare subcommunities compared to the whole bacterial communities in each sample.

**Figure S3.**
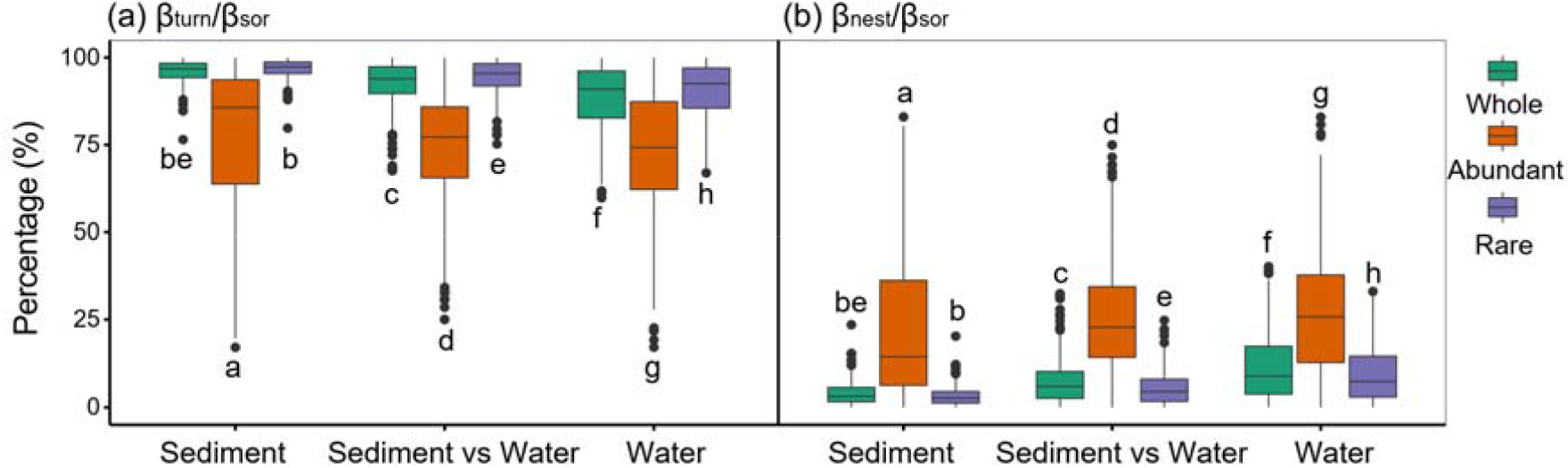
The comparison of beta-partitioning ratios between whole, abundant, and rare taxa for paired sites of only sediment samples, between sediment and water samples, and only water samples, respectively. (a) Contribution of turnover component (βturn) to total β-diversity calculated as the Sorensen dissimilarity (βsor). (b) Contribution of nestedness□resultant fraction (βnest) to total β-diversity calculated as the Sorensen dissimilarity (βsor). The different low-case letters represent significant differences of the mean values using ANOVA.

**Table S1.** The basic physicochemical properties of sediment and water samples.

|  | Minimum | Maximum | Mean | Standard Deviation | Coefficient Variation (%) |
| --- | --- | --- | --- | --- | --- |
| Sediment |  |  |  |  |  |
| pH | 8.835 | 9.835 | 9.354 | 0.246 | 2.631 |
| Conductivity (us/cm) | 204 | 1857 | 513 | 382 | 74 |
| SOC (g/kg) | 4.672 | 47.236 | 14.620 | 10.367 | 70.906 |
| TN (g/kg) | 0.280 | 2.550 | 1.092 | 0.645 | 59.045 |
| TP (g/kg) | 0.151 | 0.562 | 0.372 | 0.085 | 22.852 |
| Water |  |  |  |  |  |
| pH | 7.580 | 9.670 | 8.803 | 0.515 | 5.856 |
| Conductivity (us/cm) | 312 | 4849 | 1732 | 1416 | 82 |
| DOC (mg/L) | 1.590 | 43.850 | 16.477 | 10.438 | 63.347 |
| TN (mg/L) | 0.181 | 1.099 | 0.607 | 0.214 | 35.260 |
| TP (mg/L) | 0.039 | 0.162 | 0.100 | 0.028 | 27.871 |

**Table S2.**
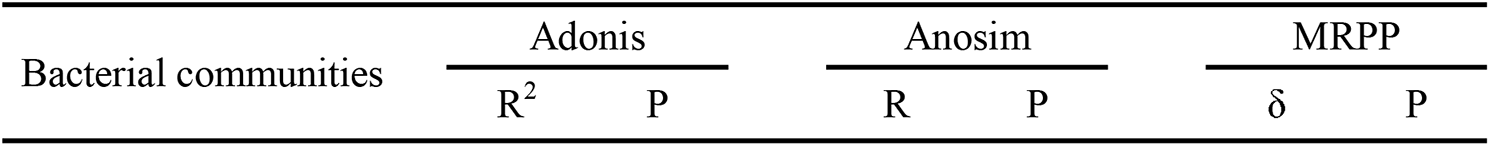
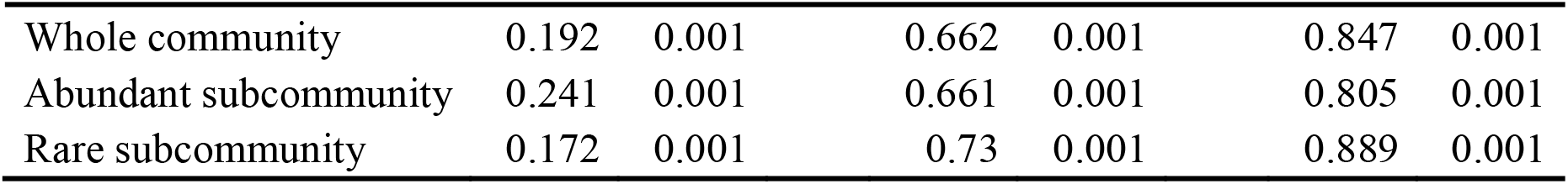
The differences between bacterial communities in sediment and water tested by analyses of Adonis, Anosim, and MRPP.

| Bacterial communities | Adonis |  | Anosim |  | MRPP |  |
| --- | --- | --- | --- | --- | --- | --- |
| | R<sup>2</sup> | P | R | P | $\delta$ | P |
| Whole community | 0.192 | 0.001 | 0.662 | 0.001 | 0.847 | 0.001 |
| Abundant subcommunity | 0.241 | 0.001 | 0.661 | 0.001 | 0.805 | 0.001 |
| Rare subcommunity | 0.172 | 0.001 | 0.73 | 0.001 | 0.889 | 0.001 |

## References

Baselga, A. (2010). Partitioning the turnover and nestedness components of beta diversity. Global Ecol. Biogeogr. 19, 134–143. doi: 10.1111/j.1466-8238.2009.00490.x

Baselga, A. (2012). The relationship between species replacement, dissimilarity derived from nestedness, and nestedness: Species replacement and nestedness. Global Ecol. Biogeogr. 21, 1223–1232. doi: 10.1111/j.1466-8238.2011.00756.x

Baselga, A., and Orme, C.D.L. (2012). betapart: an R package for the study of beta diversity. Methods Ecol. Evol. 3, 808–812. doi:

Biskaborn, B.K., Smith, S.L., Noetzli, J., Matthes, H., Vieira, G., and Streletskiy, D.A., et al. (2019). Permafrost is warming at a global scale. Nat. Commun. 10, 264. doi: 10.1038/s41467-018-08240-4

Bolger, A.M., Lohse, M., and Usadel, B. (2014). Trimmomatic: a flexible trimmer for Illumina sequence data. Bioinformatics 30, 2114–2120. doi: 10.1093/bioinformatics/btu170

Bowden, W.B. (2010). Climate change in the Arctic-permafrost, thermokarst, and why they matter to the non-Arctic world. Geography compass 4, 1553–1566. doi: 10.1111/j.1749-8198.2010.00390.x

Campbell, B.J., Yu, L., Heidelberg, J.F., and Kirchman, D.L. (2011). Activity of abundant and rare bacteria in a coastal ocean. Proceedings of the National Academy of Sciences 108, 12776–12781. doi: 10.1073/pnas.1101405108

Caporaso, J.G., Kuczynski, J., Stombaugh, J., Bittinger, K., Bushman, F.D., and Costello, E.K., et al. (2010). QIIME allows analysis of high-throughput community sequencing data. Nat. Methods 7, 335–336. doi: 10.1038/nmeth.f.303

Carbonero, F., Oakley, B.B., and Purdy, K.J. (2014). Metabolic flexibility as a major predictor of spatial distribution in microbial communities. PLoS One 9, e85105. doi: 10.1371/journal.pone.0085105

Carter, J.L., Topping, B.R., Kuwabara, J.S., Balistrieri, L.S., Woods, P.F., and Berelson, W.M., et al. (2003). Importance of sediment-water interactions in Coeur d’Alene Lake, Idaho: Management Implications. Environ. Manage. 32, 348–359. doi: 10.1007/s00267-003-0020-7

Chin, K.S., Lento, J., Culp, J.M., Lacelle, D., and Kokelj, S.V. (2016). Permafrost thaw and intense thermokarst activity decreases abundance of stream benthic macroinvertebrates. Global Change Biol. 22, 2715–2728. doi: 10.1111/gcb.13225

de Jong, A., In, T.Z.M., Meisel, O.H., Jetten, M., Dean, J.F., and Rasigraf, O., et al. (2018). Increases in temperature and nutrient availability positively affect methane-cycling microorganisms in Arctic thermokarst lake sediments. Environ. Microbiol. 20, 4314–4327. doi: 10.1111/1462-2920.14345

Debroas, D., Hugoni, M., and Domaizon, I. (2015). Evidence for an active rare biosphere within freshwater protists community. Mol. Ecol. 24, 1236–1247. doi: 10.1111/mec.13116

Farquharson, L.M., Mann, D.H., Grosse, G., Jones, B.M., and Romanovsky, V.E. (2016). Spatial distribution of thermokarst terrain in Arctic Alaska. Geomorphology 273, 116–133. doi: https://doi.org/10.1016/j.geomorph.2016.08.007

Fierer, N., Leff, J.W., Adams, B.J., Nielsen, U.N., Bates, S.T., and Lauber, C.L., et al. (2012). Cross-biome metagenomic analyses of soil microbial communities and their functional attributes. Proceedings of the National Academy of Sciences 109, 21390–21395. doi: 10.1073/pnas.1215210110

Fierer, N., and Jackson, R.B. (2006). The diversity and biogeography of soil bacterial communities. Proceedings of the National Academy of Sciences 103, 626–631. doi: 10.1073/pnas.0507535103

Gasol, J.M., Comerma, M., Garcia, J.C., Armengol, J., Casamayor, E.O., and Kojecka, P., et al. (2002). A transplant experiment to identify the factors controlling bacterial abundance, activity, production, and community composition in a eutrophic canyon-shaped reservoir. Limnol. Oceanogr. 47, 62–77. doi: 10.4319/lo.2002.47.1.0062

Gaston, K.J., Davies, R.G., Orme, C.D., Olson, V.A., Thomas, G.H., and Ding, T.S., et al. (2007a). Spatial turnover in the global avifauna. Proceedings of the Royal Society B: Biological Sciences 274, 1567–1574. doi: 10.1098/rspb.2007.0236

Gaston, K.J., Evans, K., and Lennon, J. (2007b). The scaling of spatial turnover: pruning the thicket. In: Scaling Biodiversity (eds Storch, D., Marquet, PM., and Brown, JH.). Cambridge University Press, Cambridge, UK. doi:

Gough, H.L., and Stahl, D.A. (2011). Microbial community structures in anoxic freshwater lake sediment along a metal contamination gradient. The ISME Journal 5, 543–558. doi: 10.1038/ismej.2010.132

Graham, D.E., Wallenstein, M.D., Vishnivetskaya, T.A., Waldrop, M.P., Phelps, T.J., and Pfiffner, S.M., et al. (2012). Microbes in thawing permafrost: the unknown variable in the climate change equation. The ISME Journal 6, 709–712. doi: 10.1038/ismej.2011.163

Hugerth, L.W., Larsson, J., Alneberg, J., Lindh, M.V., Legrand, C., and Pinhassi, J., et al. (2015). Metagenome-assembled genomes uncover a global brackish microbiome. Genome Biology 16, 279. doi: 10.1186/s13059-015-0834-7

In’T Zandt, M.H., Liebner, S., and Welte, C.U. (2020). Roles of thermokarst lakes in a warming world. Trends Microbiol. 28, 769–779. doi: 10.1016/j.tim.2020.04.002

Jiao, S., and Lu, Y. (2019). Soil pH and temperature regulate assembly processes of abundant and rare bacterial communities in agricultural ecosystems. Environ. Microbiol. 22, 1052–1065. doi: 10.1111/1462-2920.14815

Karlsson, J.M., Lyon, S.W., and Destouni, G. (2012). Thermokarst lake, hydrological flow and water balance indicators of permafrost change in Western Siberia. J. Hydrol. 464–465, 459–466. doi: https://doi.org/10.1016/j.jhydrol.2012.07.037

Kokelj, S.V., and Jorgenson, M.T. (2013). Advances in thermokarst research. Permafrost Periglac. 24, 108–119. doi: 10.1002/ppp.1779

Lee, Z.M.P., Poret-Peterson, A.T., Siefert, J.L., Kaul, D., Moustafa, A., and Allen, A.E., et al. (2017). Nutrient stoichiometry shapes microbial community structure in an evaporitic shallow pond. Front. Microbiol. 8, 949. doi: 10.3389/fmicb.2017.00949

Lefcheck, J.S. (2016). piecewiseSEM: Piecewise structural equation modelling in r for ecology, evolution, and systematics. Methods Ecol. Evol. 7, 573–579. doi:

Legendre, P. (2014). Interpreting the replacement and richness difference components of beta diversity. Global Ecol. Biogeogr. 23, 1324–1334. doi: 10.1111/geb.12207

Leprieur, F., Tedesco, P.A., Hugueny, B., Beauchard, O., Durr, H.H., and Brosse, S., et al. (2011). Partitioning global patterns of freshwater fish beta diversity reveals contrasting signatures of past climate changes. Ecol. Lett. 14, 325–334. doi: 10.1111/j.1461-0248.2011.01589.x

Liu, L., Yang, J., Yu, Z., and Wilkinson, D.M. (2015). The biogeography of abundant and rare bacterioplankton in the lakes and reservoirs of China. The ISME Journal 9, 2068–2077. doi: 10.1038/ismej.2015.29

Logares, R., Audic, S., Bass, D., Bittner, L., Boutte, C., and Christen, R., et al. (2014). Patterns of rare and abundant marine microbial eukaryotes. Curr. Biol. 24, 813–821. doi: 10.1016/j.cub.2014.02.050

Louca, S., Parfrey, L.W., and Doebeli, M. (2016). Decoupling function and taxonomy in the global ocean microbiome. Science 353, 1272–1277. doi: 10.1126/science.aaf4507

Lozupone, C.A., and Knight, R. (2007). Global patterns in bacterial diversity. Proceedings of the National Academy of Sciences 104, 11436–11440. doi: 10.1073/pnas.0611525104

Luo, J., Niu, F., Lin, Z., Liu, M., and Yin, G. (2015). Thermokarst lake changes between 1969 and 2010 in the Beilu River Basin, Qinghai-Tibet Plateau, China. Science Bulletin 60, 556–564. doi: 10.1007/s11434-015-0730-2

Lynch, M.D., and Neufeld, J.D. (2015). Ecology and exploration of the rare biosphere. Nat. Rev. Microbiol. 13, 217. doi:

Mackelprang, R., Burkert, A., Haw, M., Mahendrarajah, T., Conaway, C.H., and Douglas, T.A., et al. (2017). Microbial survival strategies in ancient permafrost: insights from metagenomics. The ISME Journal 11, 2305–2318. doi: 10.1038/ismej.2017.93

Malik, A.A., Martiny, J.B.H., Brodie, E.L., Martiny, A.C., Treseder, K.K., and Allison, S.D. (2020). Defining trait-based microbial strategies with consequences for soil carbon cycling under climate change. The ISME Journal 14, 1–9. doi: 10.1038/s41396-019-0510-0

Manasypov, R.M., Pokrovsky, O.S., Shirokova, L.S., Auda, Y., Zinner, N.S., and Vorobyev, S.N., et al. (2021). Biogeochemistry of macrophytes, sediments and porewaters in thermokarst lakes of permafrost peatlands, western Siberia. Sci. Total Environ. 763, 144201. doi: https://doi.org/10.1016/j.scitotenv.2020.144201

Nossa, C.W., Oberdorf, W.E., Yang, L., Aas, J.A., Paster, B.J., and Desantis, T.Z., et al. (2010). Design of 16S rRNA gene primers for 454 pyrosequencing of the human foregut microbiome. World J. Gastroentero. 16, 4135–4144. doi: 10.3748/wjg.v16.i33.4135

Oksanen, J., Kindt, R., Legendre, P.O., Hara, B., Stevens, M.H.H., and Oksanen, M.J., et al. (2007). The vegan package. Community Ecology Package 10, 631–637. doi:

Parker, S.R., West, R.F., Boyd, E.S., Feyhl-Buska, J., Gammons, C.H., and Johnston, T.B., et al. (2016). Biogeochemical and microbial seasonal dynamics between water column and sediment processes in a productive mountain lake: Georgetown Lake, MT, USA. JOURNAL OF GEOPHYSICAL RESEARCH-BIOGEOSCIENCES 121, 2064–2081. doi: 10.1002/2015JG003309

Pastick, N.J., Jorgenson, M.T., Goetz, S.J., Jones, B.M., Wylie, B.K., and Minsley, B.J., et al. (2019). Spatiotemporal remote sensing of ecosystem change and causation across Alaska. Global Change Biol. 25, 1171–1189. doi: 10.1111/gcb.14279

Pedros-Alio, C. (2006). Marine microbial diversity: can it be determined? Trends Microbiol. 14, 257–263. doi: 10.1016/j.tim.2006.04.007

Pedros-Alio, C. (2012). The rare bacterial biosphere. Annu. Rev. Mar. Sci. 4, 449–466. doi:

Pla-Rabes, S., Flower, R.J., Shilland, E.M., and Kreiser, A.M. (2011). Assessing microbial diversity using recent lake sediments and estimations of spatio-temporal diversity. J. Biogeogr. 38, 2033–2040. doi: 10.1111/j.1365-2699.2011.02530.x

Podani, J., and Schmera, D. (2016). Once again on the components of pairwise beta diversity. Ecol. Inform. 32, 63–68. doi: 10.1016/j.ecoinf.2016.01.002

Polishchuk, Y., Bogdanov, A., Polishchuk, V., Manasypov, R., Shirokova, L., and Kirpotin, S., et al. (2017). Size distribution, surface coverage, water, carbon, and metal storage of thermokarst lakes in the permafrost zone of the Western Siberia lowland. Water-Sui. 9, 228. doi: 10.3390/w9030228

Qiu, G., and Cheng, G. (1995). Permafrost in China: past and present. Permafrost Periglac. 6, 3–14. doi: 10.1002/ppp.3430060103

Quast, C., Pruesse, E., Yilmaz, P., Gerken, J., Schweer, T., and Yarza, P., et al. (2013). The SILVA ribosomal RNA gene database project: improved data processing and web-based tools. Nucleic Acids Res. 41, 590–596. doi: 10.1093/nar/gks1219

R Core Team. (2017). R: A language and environment for statistical computing, R Foundation for Statistical Computing, Vienna, Austria. https://www.R-project.org. doi:

Ren, Z., Niu, D.C., Ma, P.P., Wang, Y., Wang, Z.M., and Fu, H., et al. (2020). Bacterial communities in stream biofilms in a degrading grassland watershed on the Qinghai-Tibet Plateau. Front. Microbiol. 11, e1021. doi:

Ren, Z., Qu, X.D., Peng, W.Q., Yu, Y., and Zhang, M. (2019). Nutrients drive the structures of bacterial communities in sediments and surface waters in the river-lake system of Poyang Lake. Water-Sui. 11, e930. doi:

Ren, Z., Wang, Z.M., Wang, Y., Ma, P.P., Niu, D.C., and Fu, H., et al. (2021). Soil bacterial communities vary with grassland degradation in the Qinghai Lake watershed. Plant Soil 460, 541–557. doi:

Reyes, F.R., and Lougheed, V.L. (2015). Rapid nutrient release from permafrost thaw in arctic aquatic ecosystems. Arctic, Antarctic, and Alpine Research 47, 35–48. doi: 10.1657/AAAR0013-099

Reyon, D., Tsai, S.Q., Khayter, C., Foden, J.A., Sander, J.D., and Joung, J.K. (2012). FLASH assembly of TALENs for high-throughput genome editing. Nat. Biotechnol. 30, 460–465. doi: 10.1038/nbt.2170

Roeske, K., Sachse, R., Scheerer, C., and Roeske, I. (2012). Microbial diversity and composition of the sediment in the drinking water reservoir Saidenbach (Saxonia, Germany). Syst. Appl. Microbiol. 35, 35–44. doi: 10.1016/j.syapm.2011.09.002

Serikova, S., Pokrovsky, O.S., Laudon, H., Krickov, I.V., Lim, A.G., and Manasypov, R.M., et al. (2019). High carbon emissions from thermokarst lakes of Western Siberia. Nat. Commun. 10, 1552. doi: 10.1038/s41467-019-09592-1

Shirokova, L.S., Pokrovsky, O.S., Kirpotin, S.N., Desmukh, C., Pokrovsky, B.G., and Audry, S., et al. (2013). Biogeochemistry of organic carbon, CO2, CH4, and trace elements in thermokarst water bodies in discontinuous permafrost zones of Western Siberia. Biogeochemistry 113, 573–593. doi: 10.1007/s10533-012-9790-4

Shu, D., Guo, Y., Zhang, B., Zhang, C., Van Nostrand, J.D., and Lin, Y., et al. (2021). Rare prokaryotic sub-communities dominate the complexity of ecological networks and soil multinutrient cycling during long-term secondary succession in China’s Loess Plateau. Sci. Total Environ. 774, 145737. doi: 10.1016/j.scitotenv.2021.145737

Simek, K., Hornak, K., Jezbera, J., Nedoma, J., Znachor, P., and Hejzlar, J., et al. (2008). Spatio-temporal patterns of bacterioplankton production and community composition related to phytoplankton composition and protistan bacterivory in a dam reservoir. Aquat. Microb. Ecol. 51, 249–262. doi: 10.3354/ame01193

Torsvik, V., Ovreas, L., and Thingstad, T.F. (2002). Prokaryotic diversity - Magnitude, dynamics, and controlling factors. Science 296, 1064–1066. doi: 10.1126/science.1071698

Tranvik, L.J. (1989). Bacterioplankton growth, grazing mortality and quantitative relationship to primary production in a humic and a clearwater lake. J. Plankton Res. 11, 985–1000. doi: 10.1093/plankt/11.5.985

Viana, D.S., Figuerola, J., Schwenk, K., Manca, M., Hobæk, A., and Mjelde, M., et al. (2016). Assembly mechanisms determining high species turnover in aquatic communities over regional and continental scales. Ecography 39, 281–288. doi:

Vincent, W.F., Lemay, M., Allard, M., and Wolfe, B.B. (2013). Adapting to permafrost change: A science framework. Eos, Transactions American Geophysical Union 94, 373–375. doi: 10.1002/2013EO420002

Vucic, J.M., Gray, D.K., Cohen, R.S., Syed, M., Murdoch, A.D., and Sharma, S. (2020). Changes in water quality related to permafrost thaw may significantly impact zooplankton in small Arctic lakes. Ecol. Appl. 30, e02186. doi: 10.1002/eap.2186

Walter, K.M., Zimov, S.A., Chanton, J.P., Verbyla, D., and Chapin, F.S. (2006). Methane bubbling from Siberian thaw lakes as a positive feedback to climate warming. Nature 443, 71–75. doi: 10.1038/nature05040

Whittaker, R.J., and Fernandez-Palacios, J.M. (2007). Island biogeography: ecology, evolution, and conservation, 2nd edn. Oxford University Press, Oxford. doi:

Wu, W., Logares, R., Huang, B., and Hsieh, C.H. (2016). Abundant and rare picoeukaryotic sub-communities present contrasting patterns in the epipelagic waters of marginal seas in the northwestern Pacific Ocean. Environ. Microbiol. 19, 287–300. doi: 10.1111/1462-2920.13606

Xing, P., Guo, L., Tian, W., and Wu, Q.L. (2011). Novel Clostridium populations involved in the anaerobic degradation of Microcystis blooms. The ISME Journal 5, 792–800. doi: 10.1038/ismej.2010.176

Xue, M., Guo, Z., Gu, X., Gao, H., Weng, S., and Zhou, J., et al. (2020). Rare rather than abundant microbial communities drive the effects of long-term greenhouse cultivation on ecosystem functions in subtropical agricultural soils. Sci. Total Environ. 706, e136004. doi: 10.1016/j.scitotenv.2019.136004

Xue, Y., Chen, H., Yang, J.R., Liu, M., Huang, B., and Yang, J. (2018). Distinct patterns and processes of abundant and rare eukaryotic plankton communities following a reservoir cyanobacterial bloom. The ISME Journal 12, e2263. doi: 10.1038/s41396-018-0159-0

Yang, J., Ma, L., Jiang, H., Wu, G., and Dong, H. (2016). Salinity shapes microbial diversity and community structure in surface sediments of the Qinghai-Tibetan Lakes. Sci. Rep.-UK 6, e25078. doi: 10.1038/srep25078

Zhang, G., Yao, T., Piao, S., Bolch, T., Xie, H., and Chen, D., et al. (2017). Extensive and drastically different alpine lake changes on Asia’s high plateaus during the past four decades. Geophys. Res. Lett. 44, 252–260. doi:

Zhang, T., Barry, R.G., Knowles, K., Heginbottom, J.A., and Brown, J. (1999). Statistics and characteristics of permafrost and ground-ice distribution in the Northern Hemisphere. Polar Geography 23, 132–154. doi:

Zhao, D., Cao, X., Huang, R., Zeng, J., Wu, Q.L., and Li, M. (2017). Variation of bacterial communities in water and sediments during the decomposition of Microcystis biomass. PLoS One 12, e0176397. doi: 10.1371/journal.pone.0176397

Zou, D., Zhao, L., Sheng, Y., Chen, J., Hu, G., and Wu, T., et al. (2017). A new map of permafrost distribution on the Tibetan Plateau. Cryosphere 11, 2527–2542. doi: 10.5194/tc-11-2527-2017

